# Substrate-enhanced filamentation of 3-methylcrotonyl-CoA carboxylase in *Legionella pneumophila*

**DOI:** 10.64898/2025.12.22.694428

**Authors:** Radha P. Somarathne, Riti Shrestha, Mishghan Zehra, Cole Brodeur, David Bhella, Wing-Cheung Lai, Amit Meir, Clarissa L. Durie

## Abstract

3-Methylcrotonyl-CoA carboxylase (MCC) is a biotin-dependent carboxylase that metabolizes the amino acid leucine. MCC is present in bacteria, fungi, plants, and animals. In humans, its overexpression is linked to cancer, and its deficiency is linked to inborn errors of metabolism with severe consequences, so understanding its structure and function has far reaching implications. Here, we explore the MCC from *Legionella pneumophila*, a pathogenic bacterium with a biphasic life cycle. Our endogenous holoenzyme yielded the highest resolution cryo-EM structure of MCC to date, allowing for identification of protein components by the machine learning tool ModelAngelo, confirmed independently by mass spectrometry. We also observed, for the first time, enhanced filamentation of MCC upon substrate binding. We propose that this filamentation, previously observed in the eukaryotes, but not in bacteria, may be important for cellular processes such as differentiation of life cycle or cell division.

## Introduction

Biotin-dependent carboxylases form a conserved enzyme family that catalyzes essential reactions in carbon metabolism across all domains of life.^1–3^ 3-Methylcrotonyl-CoA carboxylase (MCC) is a biotin-dependent enzyme that catalyzes a key reaction in the metabolism of leucine, central to energy metabolism and cellular signaling.^4–6^ MCC carries out the ATP-driven carboxylation of 3-methylcrotonyl-CoA (MCoA) to form 3-methylglutaconyl-CoA, an intermediate that feeds into the tricarboxylic acid cycle via acetyl-CoA.^7^ Through this reaction, MCC links amino acid catabolism with core pathways of carbon and energy utilization. In humans, elevated MCC expression has been associated with metabolic reprogramming in several cancers.^8^ Conversely, loss of MCC activity causes 3-methylcrotonylglycinuria, one of the most frequently diagnosed deficiencies in newborns, characterized by seizures, metabolic acidosis, and progressive neurological impairment.^9–12^ MCC is also present in some, but not all, bacteria, meaning the study of MCC in bacteria allows for the discovery of fundamental biochemistry of metabolism with potential far reaching applications. Given its central role in metabolism and disease, understanding the molecular organization of MCC is essential for explaining how its structure supports catalytic function.

In general, MCC is a complex composed of six β monomers forming a ring-shape hexamer, stacked between three α monomers from each side. Overall, the holoenzyme has a α₆β₆ conformation. Structural studies from several species have revealed that MCC can adopt multiple conformational states related to ligand binding and catalysis. Crystallographic and cryo-EM analyses of MCC from *Pseudomonas aeruginosa*,^13^ *Trypanosoma brucei*,^14^ and *Leishmania tarentolae*^15^ captured different MCC forms where the co-factor, biotin, and in some instances the substrate, MCoA were bound. Comparing the conformations captured in the absence and presence of MCoA reveals that the cofactor biotin, covalently bound to a conserved lysine, is positioned differently in these different states. Biotin moves from an inactive conformation in the absence of MCoA to a conformation in which biotin’s reactive nitrogen is in proximity with MCoA’s thioester carbonyl, generating a catalytically engaged state. Because the biotin carboxylase (BC) and carboxyl transferase (CT) active sites are located on different subunits, this process requires extensive motion of the biotin carboxylase carrier protein (BCCP) domain, coordinated with movements in the BC and biotin transfer (BT) domains, consistent with a “dual-swinging-domain” mechanism.^15,16^ In *Leishmania*, MCC forms filamentous arrays through α-subunit interactions that restrict BC-domain mobility, suggesting a possible mode of structural regulation.^15^ Other acyl-CoA carboxylases have been shown to form extensive filaments with various regulatory roles,^17^ but the recent *Leishmania* report is the only example in the literature of MCC forming filaments.^15^ Together, these studies established that MCC activity is closely coupled to its conformational dynamics.

Although MCC has been structurally characterized in multiple organisms,^18^ bacterial structures have come from recombinant expression in *E. coli*, whereas native MCC has only been structurally characterized in protists. These systems have provided valuable insights into ligand binding and overall architecture but do not capture how MCC behaves in its physiological environment. It has also remained unclear whether bacterial MCCs, in their native context, can form higher-order assemblies or access the conformational diversity described in eukaryotic homologs.

The intracellular pathogen *Legionella pneumophila* provides an informative system for addressing these questions. *L. pneumophila* alternates between replication within multiple hosts (alveolar macrophages and protists) and persistence in aquatic environments, adjusting its metabolism to changing nutrient availability.^19^ During both environmental and intracellular growth, the bacterium relies heavily on amino acids, particularly serine and branched-chain amino acids such as leucine, the target of MCC, as its primary carbon and energy sources.^20,21^ Enzymes involved in leucine degradation are upregulated during stationary-phase and intracellular growth, indicating a role in metabolic remodeling.^22^ Despite this, the structure and mechanism of *L. pneumophila* MCC (LpMCC) had not been examined.

Here we report the highest-resolution structure of MCC determined by cryo-EM (2.1 Å), to date. Remarkably, this enzyme was identified through a serendipitous purification, appearing as a dominant complex within a mixed *L. pneumophila* lysate without any targeted enrichment or tagging. Multiple particle populations were resolved in the holo state where no acyl-CoA substrate was bound, suggesting conformational heterogeneity. In addition to the canonical α₆β₆ complexes, we also observed extended filamentous MCC assemblies, marking the first discovery of such stacked filaments in bacteria. While we observed fewer, shorter filaments in our holo sample, filaments represent a greater fraction of particles observed in the presence of MCoA, indicating that ligand binding promotes higher-order assembly.

Together, these findings provide a structural framework for understanding MCC function and assembly in its native bacterial context. LpMCC preserves the catalytic organization characteristic of bacterial enzymes but exhibits conformational features and ligand-dependent filament formation previously observed only in eukaryotic systems. The discovery of substrate-induced MCC filaments in a bacterium expands our understanding of how this essential metabolic enzyme adapts its architecture and dynamics to different cellular environments.

## Results and Discussion

### Identification of LpMCC using complimentary methods

The use of cryo-EM resulted in the reconstruction of multiple assembly states, the first of which we will introduce is a 2.1 Å resolution map of a 240 Å long, barrel-like structure (PDB 9TPS), with a 3-fold rotational symmetry axis and a perpendicular reflection plane of symmetry (**Fig.1a, Supporting Information, Fig. S1 and S2, Table S1**). This is the highest resolution structure of this enzyme complex reported by cryo-EM to date. As our highly pure sample was the result of the enrichment of an off-target biotin-bound protein, we first employed complimentary methods to identify the proteins present in our map.

*De novo* model building via ModelAngelo^23^ determined that the hexameric core of our structure was composed of the protein product of the gene *lpg1827*, and that the cap on either end was composed of the product of the gene *lpg1829*. Mass spectrometry finger printing of the bands observed in SDS-PAGE also identified these proteins as the major components of the isolated samples **(Supporting Information, Fig. S3, Table S2)**. Thus, two independent methods generated the same protein identifications. Furthermore, in INSeq studies reported by two groups, Roy, Shames^24^ and O’Connor^25^ the two genes coding for LpMCC α and β units (*lpg1829* and *lpg1827*, respectively) were transposon-resistant, suggesting these to be house-keeping genes essential for *L. pneumophila*’s survival. This finding was reported as a negative result in the supplementary data. Despite the many transposons made, neither group was able to produce a transposon strain in those genes. Furthermore, like in many other bacteria,^7^ the β and α units of the LpMCC are encoded within a single operon, with an enoyl-CoA hydratase/isomerase (*lpg1828*), an enzyme that plays a key role in fatty acid metabolism, located between the two genes.

Because of the high level of sequence conservation among biotin-dependent carboxylases, sequence-based annotations can be unreliable. UniProt annotated the protein Lpg1829 as acetyl-CoA carboxylase (ACC) subunit A while KEGG^26^ annotated it as MCC. Both databases annotate Lpg1827 as the propionyl-CoA carboxylase (PCC) β subunit, though KEGG orthology identified MCC. ACC, PCC, and MCC are all biotin-dependent carboxylases that carry out the same two-step carboxylation reaction on CoA-esters of different lengths. Thus, an immediate question was, what acyl-CoA ester is the likely substrate of this enzyme complex? Our data show that Lpg1829 and Lpg1827 are the α and β subunits of 3-methylcrotonyl-CoA carboxylase (MCC), respectively.

Our quaternary structure bears a striking resemblance to published structures of the MCC from multiple species,^13–15,27^ **(Fig. 1a, Supporting Information, Fig. S4)**. MCC assembles into a large multimeric complex with a conserved α₆β₆ quaternary architecture.^13^ Six β subunits form a central barrel-like core capped at both ends by trimers of α subunits. Each α subunit contains a biotin carboxylase (BC) domain that activates bicarbonate in an ATP-dependent manner, a biotin carboxyl carrier protein (BCCP) domain that carries the covalently attached biotin, and a connecting biotin transfer (BT) domain that engages the β subunit **(Fig. 1b,c)**.^16^ The β subunit consists of two homologous carboxyltransferase (CT) domains, CT-N and CT-C, whose dimer interface forms the active site for MCoA-ester substrate binding and catalysis **(Fig. 1b,c)**. As in other MCCs, the BC and CT active sites are separated by nearly 80 Å,^13^ and catalytic turnover requires large-scale movement of the BCCP domain to transfer the biotin cofactor between them.^28^ In MCC, the BC domains within each α-trimer are tightly packed, forming a compact cap distinct from the more open arrangement observed in propionyl-CoA carboxylase (PCC).^13^ Having established the overall MCC architecture, we next examined how LpMCC conforms to or diverges from these canonical structural features.

**Fig. 1.**
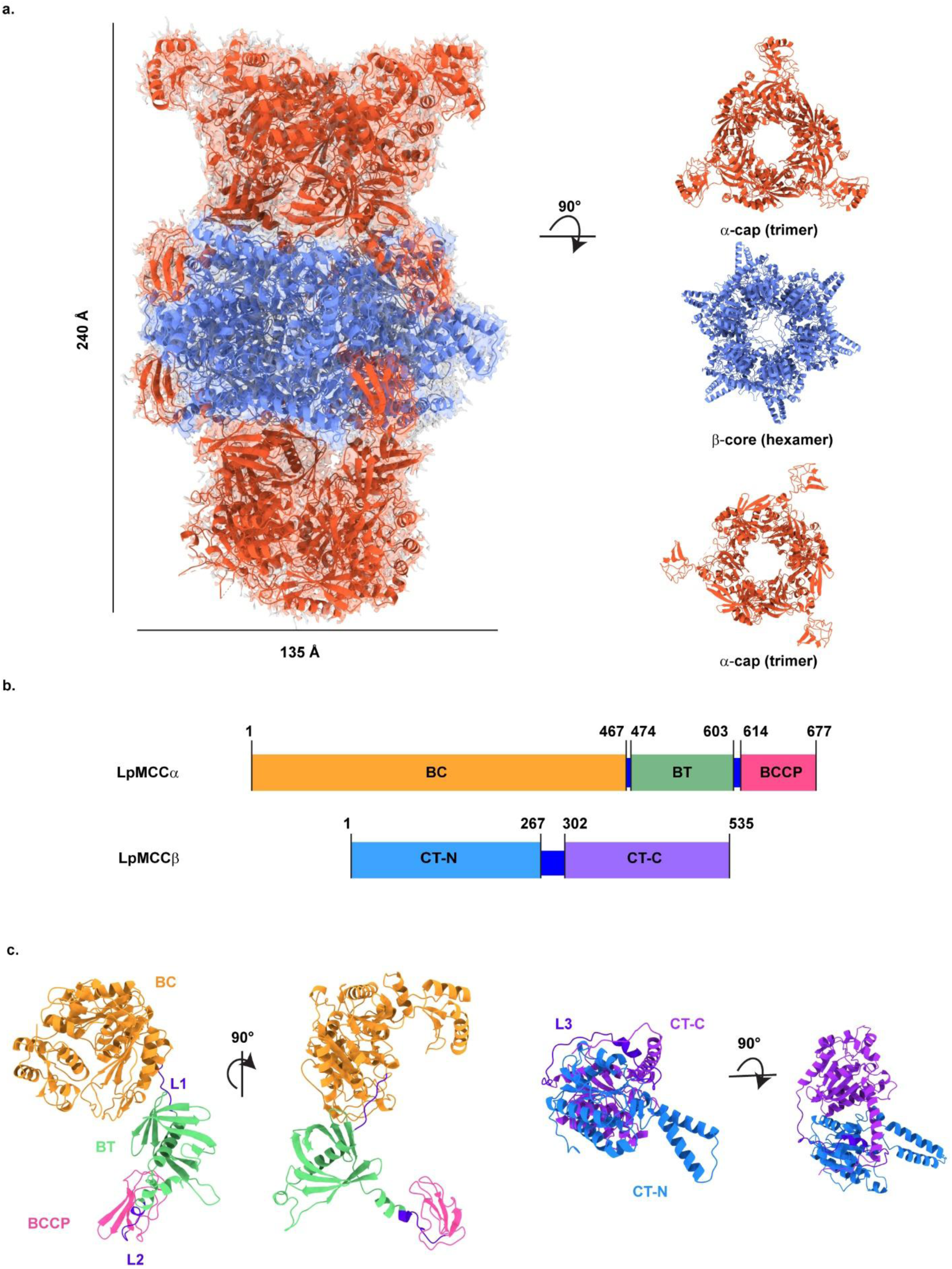
Architecture and domain organization of LpMCC. **(a)** Cryo-EM structure of the LpMCC dodecamer (PDB 9TPS) showing two α-cap trimers (red) surrounding a central β-core hexamer (blue). Right: 90° cross-sections highlighting the trimeric α-caps and hexameric β-core **(b)** Domain maps of the α- and β-subunits with residue boundaries for BC, BT, BCCP, CT-N, and CT-C domains **(c)** Structures of the α-and β-subunit protomers, colored by domain and shown in two orientations. Linkers L1, L2 and L3 which connect the domains are shown in deep blue.

### LpMCC Mirrors Established MCC Architecture and Biotin Engagement

Comparing this general description to our structure presented here, key features indicate structural homology with MCCs reported from all kingdoms of life. First, the three α subunits that make up the trimer caps are in contact, which is a feature observed in MCCs but not PCCs (**Supporting Information, Fig. S4)**.^16^ Second, the positioning of the structural elements of the biotin transfer (BT) domain of the α subunit in which an α-helix is inserted into a β-barrel is characteristic of the MCC α subunit **(Fig. 2a)** This structural feature has been reported previously, with slight variations. The *P. aeruginosa* MCC BT has a long helix inserted into an 8-stranded β-barrel while the *L. tarentolae* MCC has long helix inserted into a 7 stranded β-barrel with an additional short α-helix separating two loops or hooks that interact with the MCC β subunit. The *L. pneumophila* MCC BT displays features of both, with an 8 stranded β-barrel and a short helix with two hooks at the β subunit interface. Finally, the α₆β₆ structure presented here is homologous to the state of the human enzyme presented by Su et al described as MCC^D^.^16^ In this MCC^D^ conformation, each of the BCCP domains from each of the α subunits is associated with the CT domain of a β subunit.^16^ This conformation of the BCCP domain is “down” relative to the trimeric cap of the other domains of the α subunit.^16^

**Fig. 2.**
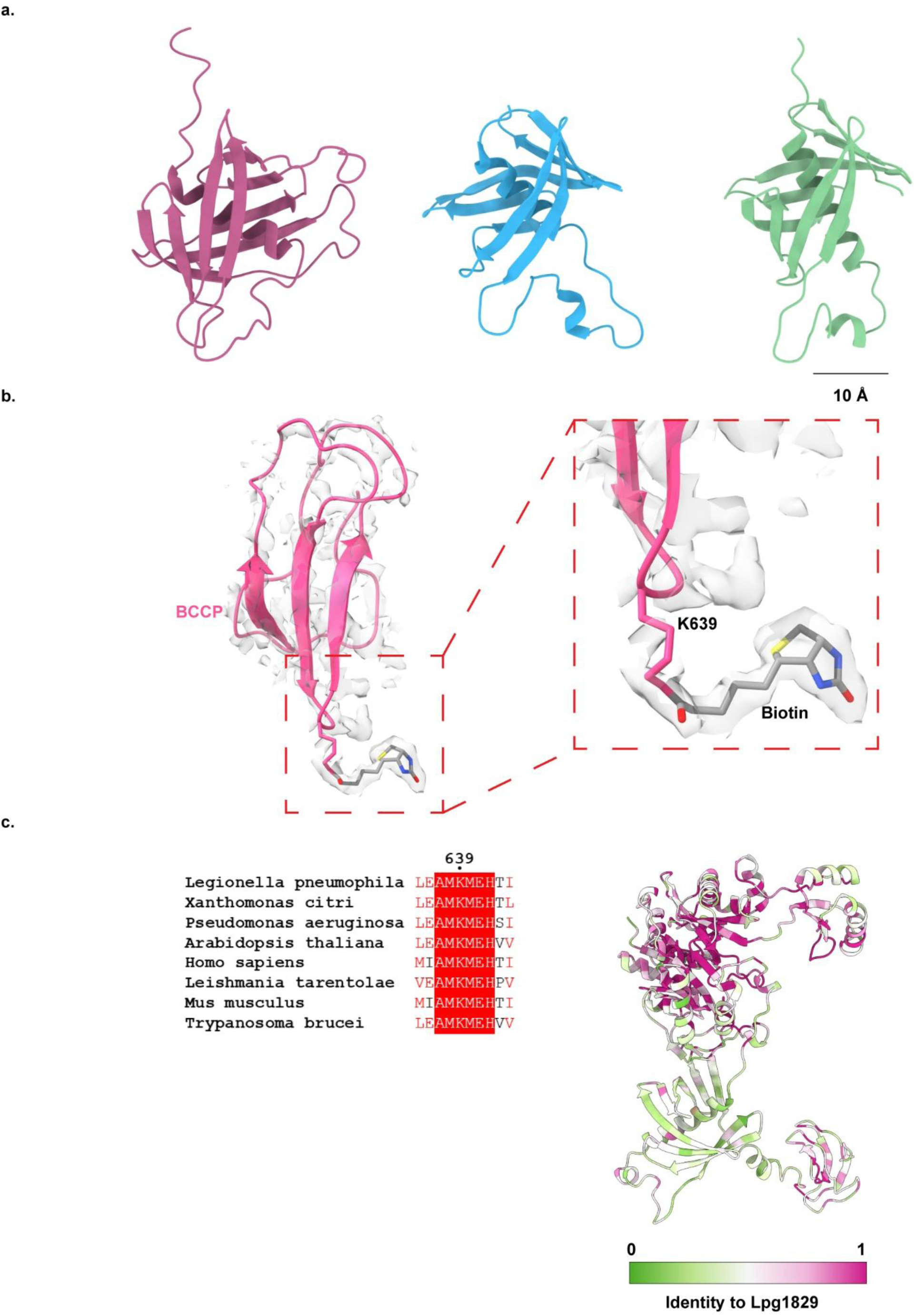
Structure and conservation of the LpMCC α subunit. **(a)** Representative BT domain folds from three species (left to right; *L. tarentolae* (PDB 8F3D), *P. aeruginosa* (PDB 3U9S), *L. pneumophila* (PDB 9TPS), shown for comparison of overall BT topology. **(b)** Cryo-EM density and model of the LpMCC BCCP domain, highlighting the covalently attached biotin at K639. **(c)** Sequence alignment of the conserved AMK motif surrounding the biotinylated lysine across diverse species, and structural mapping of residue conservation onto the LpMCC a subunit (green = low identity; pink = high identity).

We observed biotinylation of a lysine in the α subunit (K639) as shown in **Fig. 2b**. This biotinylated lysine (BTI in PDB 9TPS), biocytin, and the surrounding β strand are conserved in the other characterized structures of MCCs **(Fig. 2c)**. This region of the protein is highly conserved across all species whose MCC structures have been published, including bacteria, protozoa, and mammals **(Supporting Information, Fig. S5)**. The percent identity of Lpg1829 compared with other MCC α subunits is between 40-45%, consistent with functional homology.^29^ Because the BCCP domain is in the “down” conformation, the biocytin in the α subunit perturbs the biotin binding site in the β subunit. The *P. aeruginosa* MCC structure also revealed biotin bound in the corresponding position.^13^

### Variable α-Cap Assemblies Reveal LpMCC Heterogeneity

The β_6_ core, a hexamer formed of two stacked trimers, was observed in all states of the MCC present in our sample. We observed states with both caps, as described above, as well as states with only one cap as has been reported previously,^27^ and some states with no well-ordered trimeric cap but a partial cap on one end **(Fig. 3)**. The distribution of these populations is approximately 56% α₆β₆, 26% α_3_β_6_ (PDB 9TQC) and 18% α_1_β_6_ (PDB 9TQG). We observed flexibility in the dodecamer, particularly in the trimer caps, when subjecting the complexes to 3D Flex analysis **(Movie 1)**. The BC domains display a range of conformations with slight variations in rotation about the rotational axis, as well as slight and asymmetric contractions of the central pore between the α subunits at the apical end of the trimeric cap. The three subunits that make up the α trimeric cap stay in contact in all observed conformations, throughout interpreted motion, as reported in the recent study of the MCC from *T. brucei*.^14^

**Fig. 3.**
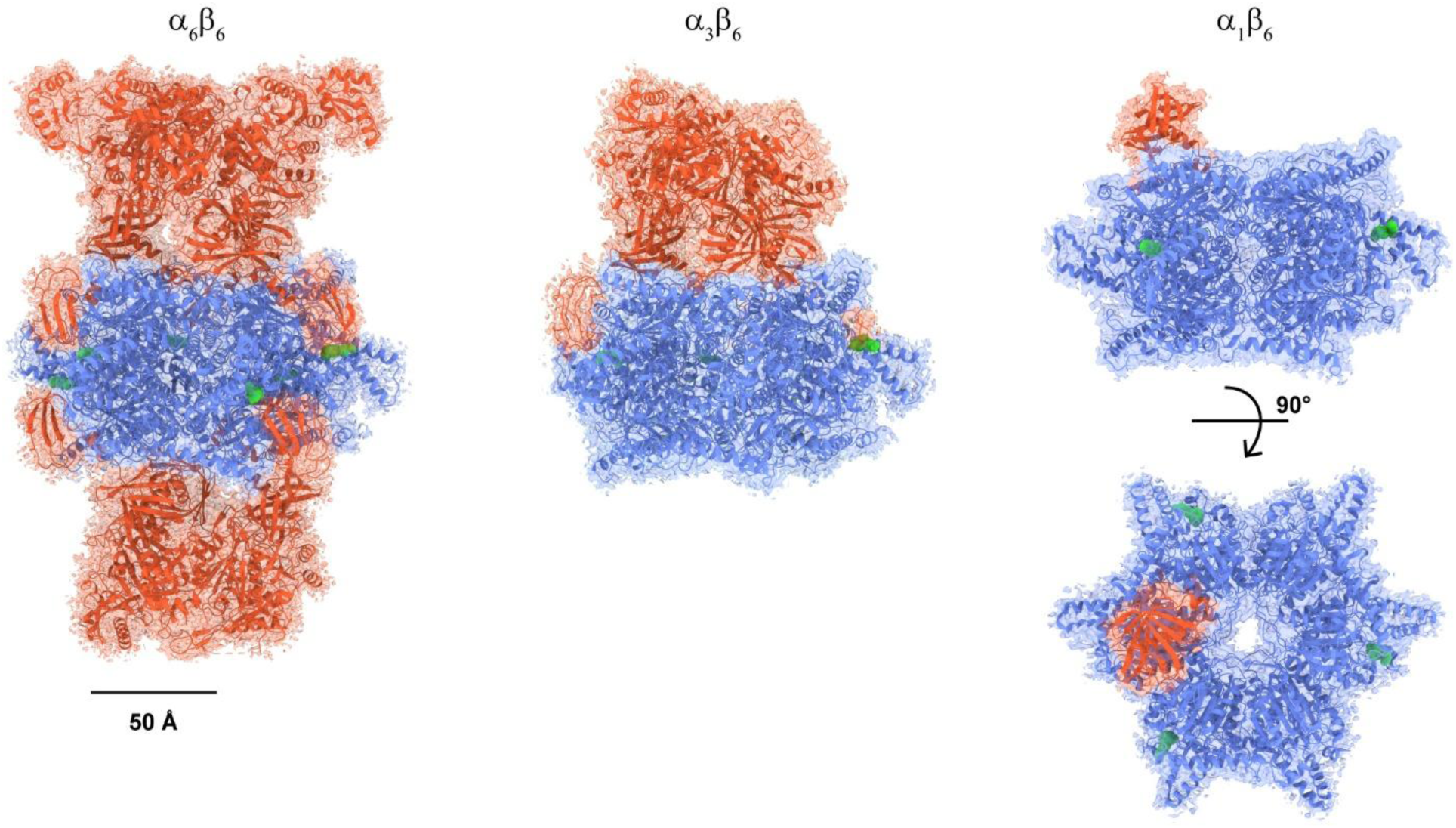
Heterogeneous α–β assembly states of LpMCC. Three predominant particle populations detected in the dataset: the complete α₆β₆ complex (left), a partially assembled α₃β₆ state (middle), (PDB 9TQC), and a minimal α₁β₆ state, (PDB 9TQG) (right). β subunits are shown in blue and α subunits in red, and biotin densities in green.

At first glance, for the α₁β₆ structure, it appeared that the α subunit caps were not associated with the β₆ particle observed here, however the 2.7 Å reconstruction revealed that one copy of the helix-in-beta-barrel motif of the BT domain of an MCC α subunit is docked into the MCC β₆ core. The remaining domains of the α subunits were not well-ordered. It is tempting to speculate that this BT domain docking could indicate an assembly pathway for the MCC complex; however, it is equally possible that this observed conformation is on a dissociation or disassembly pathway. Furthermore, a very clear biotin density, for all 6 biotin molecules, was identified in LpMCC holoenzyme α_6_β₆, α₃β₆, and the LpMCC-MCoA complex. In α_1_β₆, the biotin density was less defined, and we identified biotin in three out of the six binding sites. This was perhaps due to high flexibility of the BCCP domains. Additional experiments are required to assess the order in which these conformations appear.

Another surprising observation was the presence of filamentous assemblies made with two stacked α₆β₆ particles in length. Longer filaments have been reported in the MCC samples from *L. tarentolae,* a pathogenic protozoan, but this is the first such report from bacteria.^15^

### MCC binding to 3-methylcrotonyl-CoA induces conformational changes

To directly test the substrate specificity of our sample, we incubated MCC with the three possible CoA-ester substrates: 3-methylcrotonyl-CoA (MCoA), propionyl-CoA (PCoA), and acetyl-CoA (ACoA) and determined the structures of these samples. Previous work on other CoA-ester carboxylases have shown this to be an effective method for producing substrate-bound structures.^16^ As expected, the sample treated with MCoA yielded α₆β₆ particles with 2.3 Å resolution (PDB 9TQ5) and six molecules of 3-methylcrotonyl-CoA bound in the expected locations **(Fig. 4a and b, Supporting Information Fig. S6)**. The six MCoA molecules observed in the α₆β₆ reconstruction are bound within pockets formed at the interfaces between neighboring β subunits **(Fig. 4b)**. In each case, residues from two β monomers contribute to substrate coordination, such that each MCoA molecule is embedded between adjacent β subunits rather than being associated with a single β monomer. We also observe that the helix-turn-helix motif (residues 447 – 489) shifts in the MCoA bound structure compared to the CoA-ester-free structure to tuck in towards the core of the complex, at an angle of ∼16 ° **(Fig. 4c, Movie 2)**. We did not observe CoA-ester bound to the α₆β₆ particles reconstructed from the samples treated with PCoA or ACoA, despite achieving sufficient resolution to observe such density, if present (**Supporting Information Fig. S7**). Unlike *L. pneumophila* which bound only MCoA human MCC has been shown to bind MCoA, PCoA, and ACoA. Notably, human MCC showed a similar closing of the helix-turn-helix motif upon CoA-ester binding only for MCoA and ACoA, but not PCoA.^16,27^ Notably, while the helix-turn-helix structural motif is observed in all experimentally determined MCC structures, and while the MCC β subunits have higher percent identity than the α subunits **(Fig. 4d, Supporting Information Fig. S8)**, this motif which is associated with substrate binding is very poorly conserved across species.

**Fig. 4.**
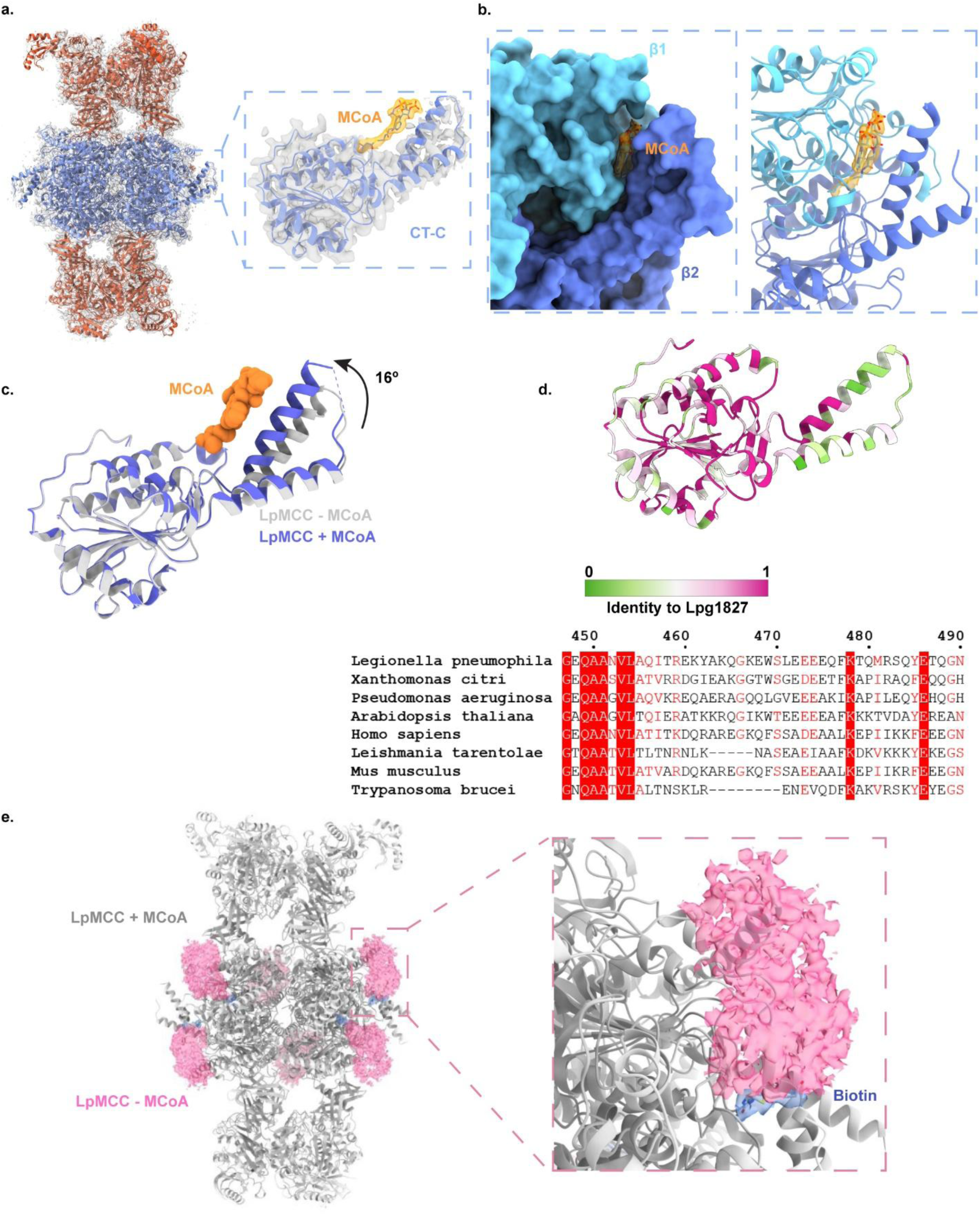
Structure of LpMCC in the presence of 3-methylcrotonyl-CoA. **(a)** Cryo-EM reconstruction of MCoA-bound LpMCC (PDB 9TQ5), showing an α₆β₆ holoenzyme with substrate bound at all six CT-C active sites. Right: zoomed view of the CT-C domain with MCoA density. **(b)** Surface representation of the β subunit CT-C domain showing the MCoA binding pocket, with bound MCoA shown as sticks. **(c)** Superposition of the apo (gray) and MCoA-bound (blue) LpMCC CT-C domains highlighting movement at the helix–turn–helix motif upon MCoA binding. **(d)** Sequence alignment across MCC homologs showing low sequence conservation within the helix–turn–helix region. Top: structural identity mapping onto the LpMCC β subunit; bottom: multiple sequence alignment of residues 450–490. **(e)** Comparison of holomap with no CoA-ester (pink) and MCoA-bound LpMCC model (grey) showing loss of ordered density for the BCCP domain.

There were additional unanticipated findings in the 3-methylcrotonyl-CoA dataset. In the 3-methylcrotonyl-CoA–bound reconstruction, density corresponding to the BCCP domain is markedly weakened and only becomes apparent at lower contour thresholds **(Fig. 4e)**, suggesting that this domain is more flexible or disordered compared with the CoA-ester–unbound state. Another possibility is that binding of the MCoA substrate induces a conformational change that repositions the biotin moiety, thereby promoting delocalization of the BCCP domain **(Supporting Information Fig. S7)**.

### Binding of 3-methylcrotonyl-CoA enhances filamentation

Finally, the presence of filaments increases in both frequency and length upon substrate binding, with assemblies extending beyond three α₆β₆ units, approximately twice the length of those observed in the absence of CoA-ester substrate **(Figure 5a**, **Table 1, Supporting Information Fig. S6)**. While substrates, inhibitors, and other compounds have been reported to influence filament formation in other biotin dependent carboxylases^15,30^, substrate associated changes in MCC filamentation have not been previously described.

**Fig. 5.**
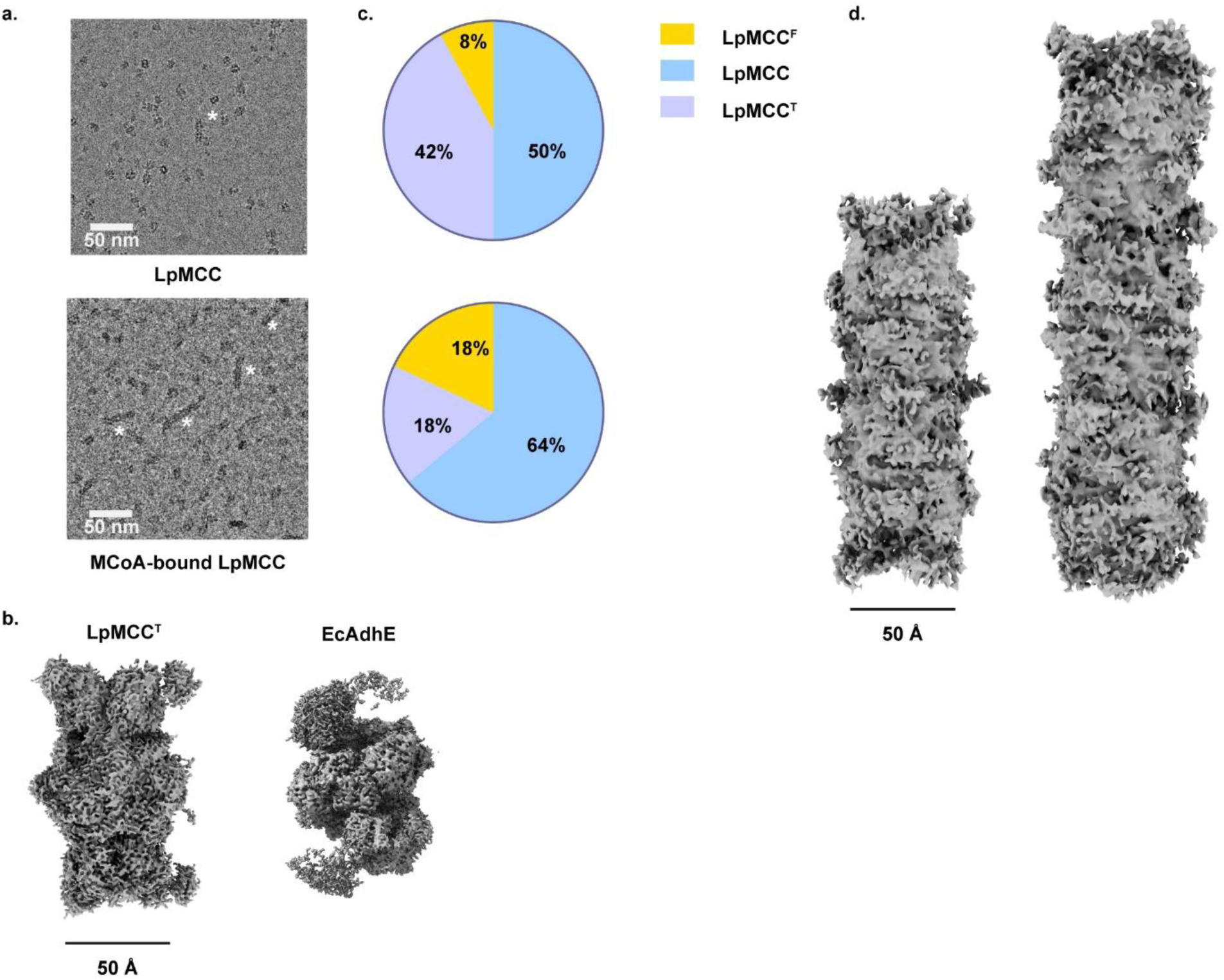
Distribution of LpMCC particle assemblies in the absence and presence of MCoA. **(a)** Representative cryo-EM micrographs of LpMCC in the absence of MCoA (top) and in the presence of MCoA (bottom). Filamentous assemblies (LpMCC^F^) are indicated by asterisks (*). **(b)** From left to right, 3D reconstructions of a tilted particle (LpMCC^T^) from this study and a subunit of the filament or spirosome of *Escherichia coli* AdhE (PDB 6AHC, EMD-9623) from Kim et. al sowing a tilted conformation **(c)** Pie charts showing the relative proportions of LpMCC particle populations, obtained by heterogenous refinement, observed for LpMCC (top) and MCoA-bound LpMCC (bottom), where LpMCC^T^ = tilted particles and LpMCC^F^ = filaments / filamentous assemblies. **(d)** 3D reconstruction of LpMCC^F^ assemblies containing 2 and 3 LpMCC^T^ particles

**Table 1.** Frequency of LpMCC^F^ particles per 100 micrographs (N/D – Not Detected)

|  | 2 particles | 3 particles | 3+ particles |
| --- | --- | --- | --- |
| LpMCC + MCoA | 100.3 | 55.6 | 1.2 |
| LpMCC | 3.2 | N/D | N/D |

In the cryo-EM micrographs, we observe a subset of particles that are structurally distinct from the canonical α₆β₆ LpMCC holoenzyme **(Supporting Information Fig. S6)**. These particles, which we refer to as Tilted LpMCC (LpMCCᵀ), are frequently associated with Filamentous assemblies (LpMCC^F^) of varying lengths. Although these particles could initially be interpreted as α₆β₆ LpMCC viewed in an alternative orientation, particle selection and independent 3D reconstructions indicate that LpMCCᵀ represents a structurally distinct population **(Fig. 5b)**. LpMCCᵀ particles display a tilted β₆ core, resulting in a slightly elongated appearance, while the BT domain appears distorted relative to the canonical structure.

Quantification using particle counts from cryoSPARC following sorting into different conformations (heterogeneous refinement in cryoSPARC) shows that, in the absence of MCoA, LpMCC and LpMCCᵀ single particles are present at comparable levels, and filamentous assemblies represent only a minor population **(Fig. 5c)**. In contrast, in the presence of MCoA, the relative abundance of LpMCCᵀ decreases, with a corresponding increase in filamentous assemblies by approximately 48-fold, suggesting that a subset of LpMCCᵀ particles contributes to filament formation under these conditions. Neither propionyl-CoA nor acetyl-CoA enhanced filamentation (data not shown). In the absence of MCoA, the sparse filament population is composed primarily of two particles of LpMCCᵀ, whereas MCoA binding is associated with both increased filament length and an increased proportion of higher-order assemblies containing three or more LpMCCᵀ particles **(Table 1**, **Fig. 5b and d)**.

These observations indicate that filament formation in LpMCC is closely linked to the presence of 3-methylcrotonyl-CoA and involves a structurally distinct tilted particle population. In contrast to LtMCC, where filament formation has been observed in the absence of substrate and interpreted as an inactive or sequestered state, LpMCC filaments arise under substrate-bound conditions, leading to a question whether polymerization is inherently associated with enzyme inhibition. The reduction in free LpMCCᵀ particles together with an increase in filament length and abundance supports a model in which LpMCC exists in equilibrium with a tilted LpMCCᵀ conformation, and this equilibrium is shifted by MCoA binding toward states that favor higher-order assembly.

In recent years, evidence accumulate to show metabolic enzymes such as ACC,^30,31^AdhE^32,33^ **(Fig. 5b)** and CTP synthase^34^ assemble and function in filament-dependent manner in both prokaryotic to eukaryotic organisms. Metabolic enzyme filamentation is attributed to regulation, including allosteric control of enzyme activity and nutrient availability. Dynamic assembly and disassembly of filaments allows for rapid responses to changing cellular or environmental metabolic state, suggesting that it serves as a mechanism to modulate metabolic flux in response to cellular demand.^35,36^ In this regard, filament formation in bacteria has been proposed to represent an extension of the bacterial cytoskeleton, with roles in regulating cell shape, as exemplified by CTP synthase filaments.^34^ Intriguingly, our mass spectrometry analysis detected low levels of the cytoskeletal proteins MreB and FtsZ in LpMCC purifications, suggesting a potential, albeit low-abundance, association between LpMCC and components of the bacterial cytoskeleton.

Importantly, the enhancement of filamentation of MCC upon MCoA binding has not previously been reported. While the triggers for filament enhancers have been described in other metabolic enzymes (e.g. glucokinase 1),^37^ MCC’s substrate has not been described as its polymerization enhancer. Filaments have, however, been observed, notably in the MCC of *Leishmania tarentolae*.^15^ Both the bacterial pathogen *Legionella pneumophila* and the protozoan pathogen *L. tarentolae* exhibit biphasic lifecycles with markedly different nutrient availabilities depending on life cycle phase and corresponding environment. *L. pneumophila* cycles between freshwater sources and eukaryotic host cells such as amoebae or alveolar macrophages, and its virulence factors include effectors that redirect host cell machinery and nutrients, including amino acids, to the *Legionella*-containing vacuole for intracellular multiplication. *L. tarentolae* cycles between the amino acid–rich environment of the host gut and the glucose-rich environment of the host bloodstream. Thus, both organisms have evolved to thrive in environments with distinct nutrient profiles.^15^

MCC plays a central role in leucine metabolism and energy production, and its functional importance is therefore likely to vary across different phases of the pathogen lifecycle. In this context, the substrate-dependent filamentation observed in LpMCC may represent a regulatory mechanism, as has been described for other filament-forming enzymes.^17^ The tilted and distorted features of LpMCCᵀ, particularly within the BT domain, may facilitate the intermolecular contacts required for filament formation, providing a structural basis for such regulation in response to metabolic conditions.

The work presented here lays the foundation for additional studies into the linkage between metabolism and virulence. The connection between metabolism and virulence via the stringent response is well established, but it is not known whether MCC plays a role. The functional consequence of filamentation in *L. pneumophila* is yet to be explored. Filamentation could either increase or decrease MCC activity and thus carbon flux through leucine metabolism. Another open question is how MCC is regulated across the *L. pneumophila* lifecycle, whether that regulation is by transcription, translation, or sequestration/activation by filamentation. Additionally, a second biotin-dependent carboxylase was also isolated during this investigation, with characterization under way (data not shown).

## Acknowledgements

The authors acknowledge the use of the Titan Krios G4 Cryo-TEM at the Electron Microscopy Core Facility, which is supported and administered by the Office of Research, Innovation, and Impact at the University of Missouri. The authors also thank Dr. Min Su from the University of Missouri Electron Microscopy Core for technical support and training in microscopy.

The computation for this work was performed on the high-performance computing infrastructure operated by Research Support Solutions in the Division of IT at the University of Missouri, Columbia MO https://doi.org/10.32469/10355/97710

Mass Spectrometry analyses were performed by the Mass Spectrometry Technology Access Center at the McDonnell Genome Institute (MTAC@MGI) at Washington University School of Medicine, supported by the Diabetes Research Center/NIH grant P30 DK020579, Institute of Clinical and Translational Sciences/NCATS CTSA award UL1 TR002345, and Siteman Cancer Center/NCI CCSG grant P30 CA091842.

We thank Dr. Adam Yokom, Dr. Robert Kazmierczak, Dr. David Buckley (UoM) and Dr Martin Rennie and Dr Michael Capper (UoG) for helpful conversations.

Research reported in this publication was supported by the National Institute of General Medical Sciences of the National Institutes of Health under award number R35GM150663 to CLD and Wayne L. Ryan Graduate Fellowship, Ryan Foundation to MZ.

AM and DB are supported by MRC-University of Glasgow Centre for Virus Research core and the Medical Research Council Award (MC_UU_00034/1) to DB.

## Author Contributions

C.L.D., A.M., and R.P.S. conceived and designed the experiments. R.P.S., R.S., M.Z., C.B, and W-L.C. performed the experiments. M.Z. grew the cultures. R.P.S., R.S., M.Z., C.B, and W-L.C. purified the complexes and prepared samples. R.P.S. performed structural experiments and analyses for the LpMCC and MCoA-bound LpMCC samples. R.S. performed the structural experiments and analyses for the ACoA and PCoA samples. A.M. and D.B. identified the complex through ModelAngelo. A.M. performed model building, refinement with support from R.P.S., and validation. R.S. and C.B. performed the MSA analyses. R.S. coordinated the mass spectrometry experiments and analysis. D.B. provided access to modeling resources. A.M and C.L.D. directed the work. All the authors have edited the manuscript.

## Materials and Methods

### Bacterial culture and growth conditions

*Legionella pneumophila* strain Lp01 (generously provided by Craig Roy, Yale University) containing a *ΔdotB* deletion and harboring plasmid pDotBStrep, was used for all experiments. Bacteria were maintained on charcoal yeast extract (CYE) agar supplemented with streptomycin (100 μg/mL) and chloramphenicol (10 μg/mL)^38^. For liquid culture, *L. pneumophila* was grown in ACES-buffered yeast extract (AYE; pH 6.9) medium containing L-cysteine (0.4 mg/mL) and ferric nitrate (0.135 mg/mL). Solid AYE plates were prepared by adding agar (15 g/L) and activated charcoal (2 g/L)^39^. Cultures were incubated at 37 °C with shaking at 220 rpm. Protein expression was induced with 1 mM isopropyl β-D-1-thiogalactopyranoside (IPTG) at the time of inoculation, and cultures were grown to OD_600_ 3.4 - 3.6 absorbance units.

### Protein purification

Cells were harvested by centrifugation at 10,000 × g for 10 min and resuspended in freshly prepared LP buffer (50 mM Tris pH 8.0, 200 mM NaCl, 2 mM EDTA, 20 mM MgSO₄) containing 0.5 M sucrose, 0.1 mg/mL lysozyme, 0.1 mg/mL DNase I, and protease inhibitor cocktail (Roche). The suspension was rotated gently for 45 min at 4 °C, followed by centrifugation at 14,000 × g for 12 min. The cell pellet was resuspended in LP buffer. Cells were lysed French press at 14,000 psi. The lysate was clarified by centrifugation at 17,300 × g for 20 min to remove debris, followed by ultracentrifugation at 167,000 × g for 2 h. The soluble fraction was applied to a 5 mL StrepTrap XT column (Cytiva) pre-equilibrated with LP buffer and connected to a Bio-Rad NGC FPLC system. Bound proteins were eluted with LP buffer containing a linear gradient of 0–50 mM biotin. Fractions corresponding to the major absorbance peak at 280 nm were pooled and analyzed by SDS–PAGE to verify purity and molecular weight **(Supporting Information, Fig. S1).**

### Substrate binding

LpMCC samples in LP buffer were incubated on ice with 10 mM acetyl-CoA (#A2056), propionyl-CoA (#P5397), or methylcrotonyl-CoA (M3013) from Sigma Aldrich, for 1.5 h. The mixture was directly used for cryo-EM sample preparation.

### Cryo-electron microscopy

All microscopy-related sample preparation and data collection were performed at the Electron Microscopy Core at the University of Missouri.

### Sample preparation

Purified LpMCC and substrate-bound samples were prepared for cryo-electron microscopy following standard vitrification procedures. Quantifoil R1.2/1.3 200-mesh Cu grids (Electron Microscopy Sciences) were glow discharged for 15 s at 5 mA using a PELCO easiGLow (Ted Pella). A 3.5 µL aliquot of each protein sample was applied to each grid using a FEI Vitrobot Mark IV (Thermo Fisher Scientific) operated at 4 °C and 100% relative humidity.

### Data collection

Cryo-EM data for LpMCC and LpMCC-MCoA samples were collected on a FEI Titan Krios G4 transmission electron microscope (Thermo Fisher Scientific) operated at 300 kV. The microscope was equipped with a Gatan K3 direct electron detector operated in counting mode and a Gatan BioQuantum. Automated data collection was carried out using EPU software (Thermo Fisher Scientific). Micrographs were recorded at a magnification of 165,000x, corresponding to a calibrated pixel size of 0.75 Å/pixel with a defocus range of −0.8 to −1.6 µm. The total dose rate on the detector was 40 e^−^/Å^2^ with a total exposure time of 3.22 s, and 22 frames.

For LpMCC,a total of 31,641 micrographs and for LpMCC-MCoA, LpMCC-ACoA and LpMCC-PCoA 14,859, 9,011 and 11,825 micrographs were collected respectively.

### Data processing

Data processing was carried out in CryoSPARC^40^ v4.7.1 running on the University of Missouri Hellbender high-performance computing cluster. Motion correction and dose-weighting were performed using the patch-based workflow, followed by CTF estimation for each movie. Micrographs with poor CTF fits, excessive drift, or visibly thick ice were removed during manual curation. For LpMCC, 3,863 particles were initially selected by blob picking and used to generate reference 2D class averages which were then used as templates for template picking **(Supporting Information, Fig. S1)**. Template picking identified ∼21.7 million particles across the curated micrographs. After cleanup and re-extraction, ∼1.5 million particles were taken forward into multiple rounds of 2D classification to remove damaged or misaligned particles and enrich for clear secondary-structure features. A final set of 435,252 particles was retained for 3D analysis. A single-class *ab initio* reconstruction was generated and refined by homogeneous and non-uniform refinement in C1, resulting in a 2.1 Å reconstruction from the full particle set. In parallel, a three-class *ab initio* run was used to examine particle heterogeneity. Each class was refined independently using heterogeneous followed by homogeneous refinement, yielding reconstructions at 2.5 Å (114,978 particles), 2.7 Å (78,376 particles), and 2.2 Å (241,903 particles).

Processing of the LpMCC–MCoA dataset followed the same general workflow **(Supporting Information, Fig. S4)**. A total of 14,859 micrographs were retained. Multiple rounds of 2D classification separated single particles from filamentous assemblies. The single-particle subset contained 434,423 particles. This subset was used for *ab initio* reconstruction followed by heterogeneous and homogeneous refinement in C1, yielding a 2.3 Å reconstruction from 338,596 particles.

For processing filamentous assemblies, 95,891 particles identified during 2D classification were used to generate templates for particle picking. Approximately 600,000 filament-containing particles were extracted using a 2,000 px box size and downsampled to 500 px for subsequent analysis. Multiple rounds of 2D classification were performed to enrich for well-defined filament views. Tilted particles, LpMCC^T^ and filaments, LpMCC^F^, composed of two, three, or more stacked LpMCC^T^ particles were separated and quantified using heterogenous refinement and *ab initio* classification respectively. Further refinement of these filament subsets did not yield high-resolution reconstructions.

Data processing and collection parameters for other substrate bound particles (LpMCC-ACoA and LpMCC-PCoA) are summarized in **Supporting Information, Table S1.**

### Atomic model building and refinement

The Lpg1827 and Lpg1829 components were identified using the automated machine-learning tool ModelAngelo^23^, which generated *ab initio* atomic models based on the full proteome of *Legionella pneumophila* strain Philadelphia (accession: NZ_CP013742.1). The top-scoring hits corresponded to Lpg1827 and Lpg1829 (β and α subunits, respectively), which were further validated by pBLAST searches. AlphaFold3^41^ predictions of the identified proteins or specific domains were superimposed onto the fragmented chains generated by ModelAngelo and subsequently fitted into the cryo-EM density maps using ChimeraX^42^. Manual model adjustment was performed by rounds of Coot^43^ and molecular dynamics–based refinement using ISOLDE^44^, followed by real space refinement in Phenix.^45,46^

Sequence assignments were confirmed by detailed inspection against high-resolution density features. Ligands (biotin, BTI and methylcrotonyl-CoA, MCoA) were fitted into density by superimposing pre-existing structures (PDB 3U9S)^13^ onto corresponding domains, following refinement. The MCoA ligand was downloaded from (PDB 8J4Z)^16^, where it was named TW3.

To further increase confidence in the assignment of unique proteins to previously uncharacterized densities, reverse searches were conducted against known structures of MCC, ACC, and PCC, employing both DALI^47^ and AlphaFold3 predictions of candidate proteins as queries. Final models were refined in Phenix using real-space refinement optimized for cryo-EM data, and validated using its comprehensive EM validation. All molecular graphics and structural analyses were carried out in ChimeraX. Model statistics are summarized in **Supporting Information, Table S1.**

### Protein Identification by Mass Spectrometry Analysis

Excised protein gel bands **(Supporting Information, Fig. S3)** were submitted to Mass Spectrometry Technology Access Center at the McDonnell Genome Institute, Washington University School of Medicine for in-gel digestion and tandem mass spectrometry analysis. Tandem mass spectra were extracted using proprietary software (version not recorded). The resulting MS/MS data were analyzed using Mascot (version 3.1.0; Matrix Science, London, UK) against the *Legionella pneumophila* Philadelphia-1 protein database, assuming trypsin digestion. Search parameters included 10.0 ppm parent ion and 0.05 Da fragment ion tolerances; carbamidomethylation of cysteine was set as a fixed modification, and deamidation (N/Q), oxidation (M), and N-terminal acetylation as variable modifications (**Supporting Information, Table S2).** Peptide and protein identifications were validated in Scaffold (v5.3.4; Proteome Software Inc., Portland, OR) using Peptide Prophet algorithm^48^ at >90% probability and Protein Prophet algorithm^49^ at >99% probability, requiring at least two identified peptides. Proteins sharing significant peptide evidence and indistinguishable by MS/MS analysis were grouped according to the principles of parsimony.

### Sequence Alignment and Analysis

Amino acid sequences of the MCC α (biotin carboxylase and biotin carboxyl carrier protein domains) and MCC β (carboxyl transferase domain) subunits from *Legionella pneumophila* and representative bacterial and eukaryotic species were retrieved from the UniProt database. Multiple sequence alignments were generated using Clustal Omega^50^, ensuring accurate alignment of conserved and variable regions. Alignments were manually inspected and conserved residues and domain architecture were visualized using ESPript 3.0^51^, and domain boundaries were annotated based on InterPro predictions.

