## Supporting Information for "Substrate-enhanced filamentation of 3-methylcrotonyl-CoA carboxylase in *Legionella pneumophila*"

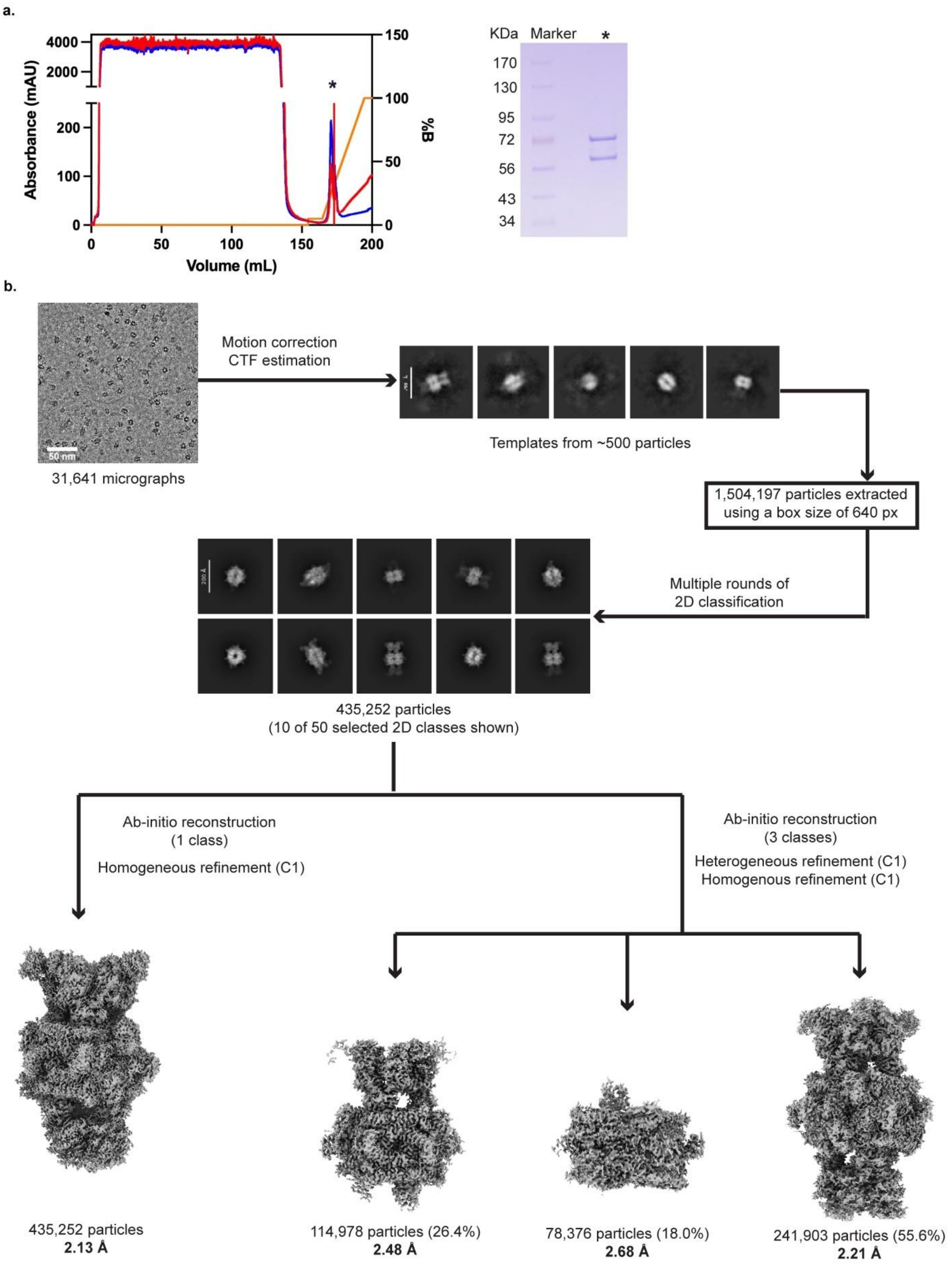


**Fig. S1 Purification and cryo-EM structure determination of *Legionella pneumophila* methylcrotonyl-CoA carboxylase (LpMCC).**

**(a)** StrepTrap affinity chromatography profile of LpMCC. The asterisk (*) marks the LpMCC elution peak; the corresponding fraction was analyzed by SDS–PAGE (right). **(b)** Cryo-EM workflow used to resolve distinct structural populations of LpMCC.

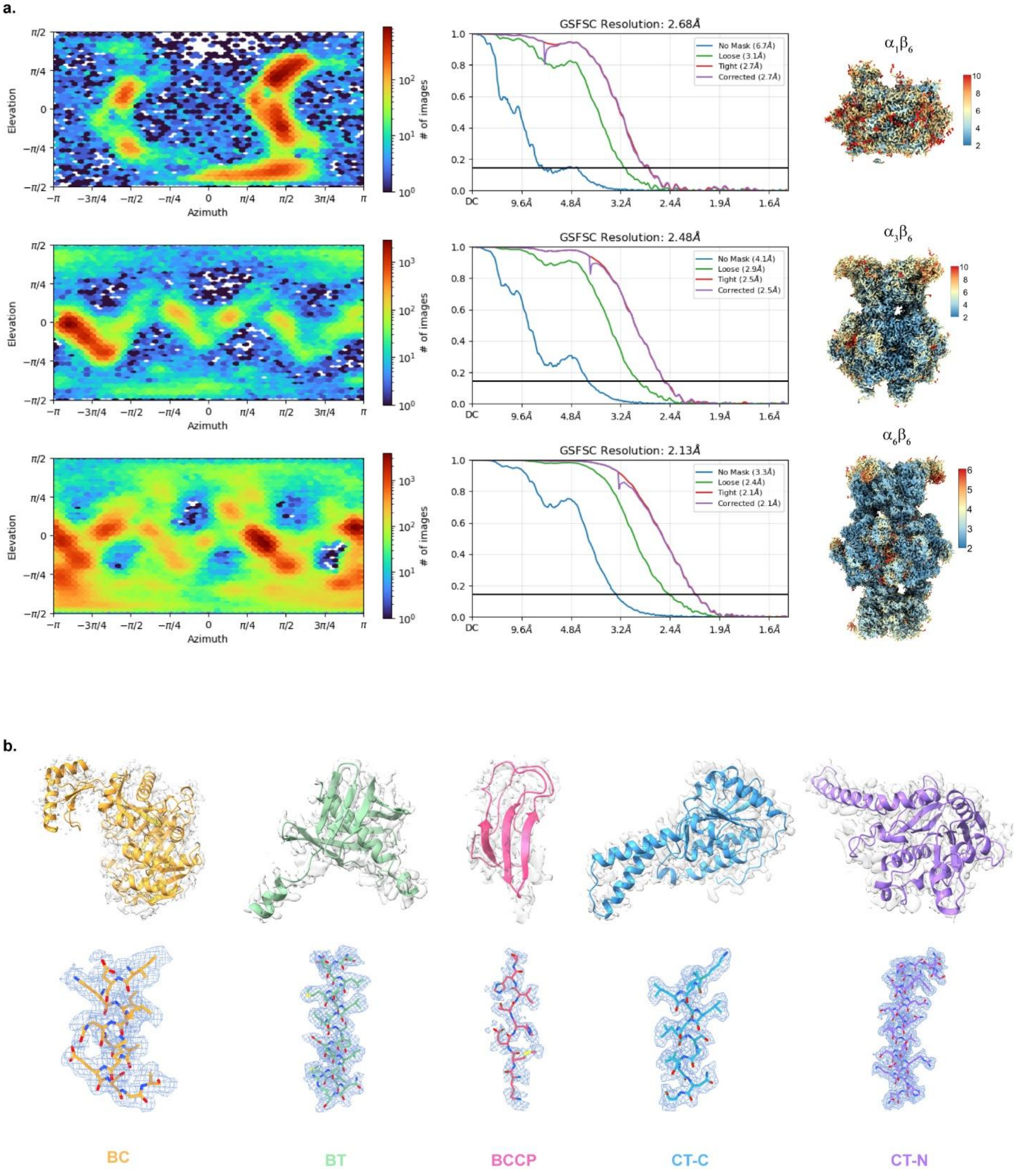


#### Fig. S2 Cryo-EM map quality and domain-level density of LpMCC (PDB 9TPS)

**(a)** Euler angle distributions and gold-standard Fourier shell correlation (GSFSC) curves for representative LpMCC reconstructions where resolutions are reported at the FSC = 0.143 criterion and local resolution maps of the refined densities, colored according to the resolution scale (Å). **(b)** Local cryo-EM density maps with fitted models highlighting domains of the α subunit (BC, BT, and BCCP) and the β subunit (CT-C and CT-N). Representative regions are shown to illustrate side-chain density.

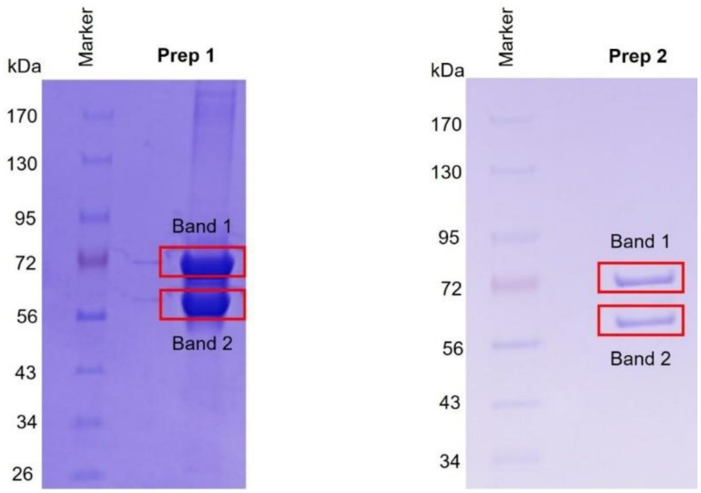


**Fig. S3 SDS–PAGE analysis of LpMCC preparations used for mass spectrometry.**

SDS–PAGE gels of two independent LpMCC preparations (Prep 1 and Prep 2) used for mass spectrometry fingerprinting. Bands selected for analysis are indicated (Band 1 and Band 2)

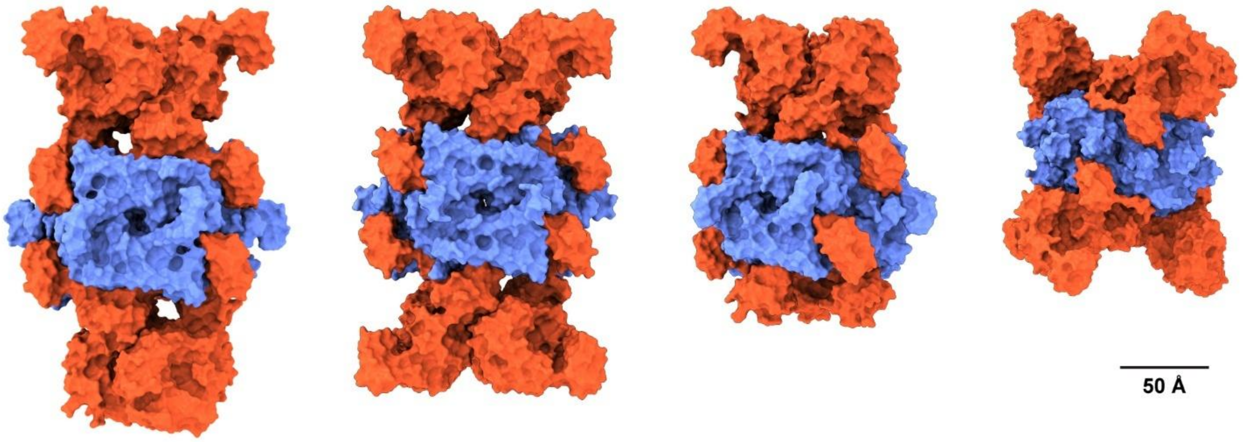


**Fig. S4 Structural comparison of biotin-dependent carboxylases.**

From left to right, structures of biotin-dependent carboxylases shown in equivalent orientations: *Legionella pneumophila* methylcrotonyl-CoA carboxylase (LpMCC) from this study (PDB 9TPS); methylcrotonyl-CoA carboxylase from *Trypanosoma brucei* (PDB 8RTH); human 3-methylcrotonyl-CoA carboxylase in the apo state (PDB 8XL6); and human propionyl-CoA carboxylase (PDB 7YBU). α subunit domains are shown in red and β subunit domains in blue.

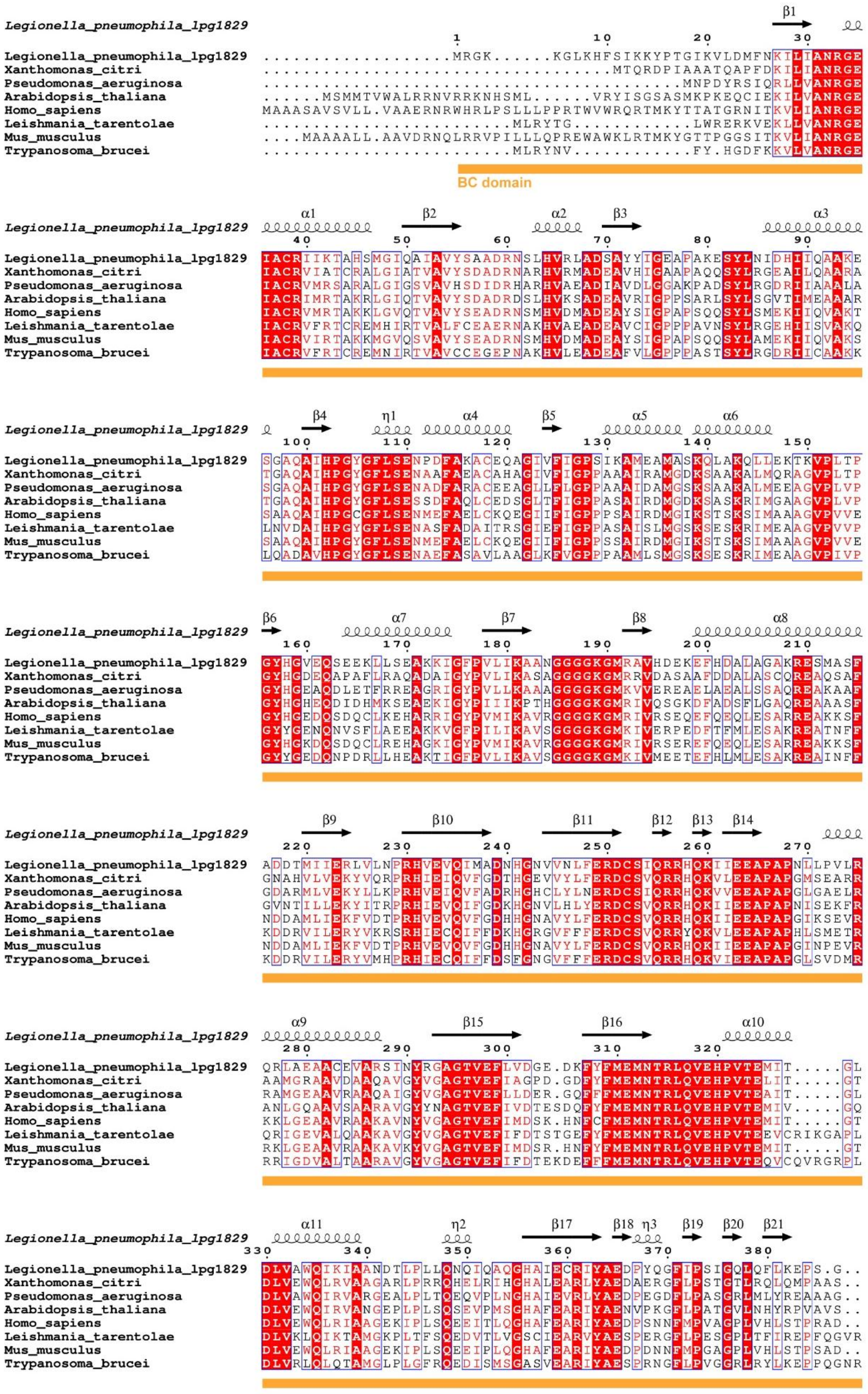

Legionella_pneumophila_lpg1829
Xanthomonas_citri
Pseudomonas_aeruginosa
Arabidopsis_thaliana
Homo_sapiens
Leishmania_tarentolae
Mus_musculus
Trypanosoma_brucei
BC domain

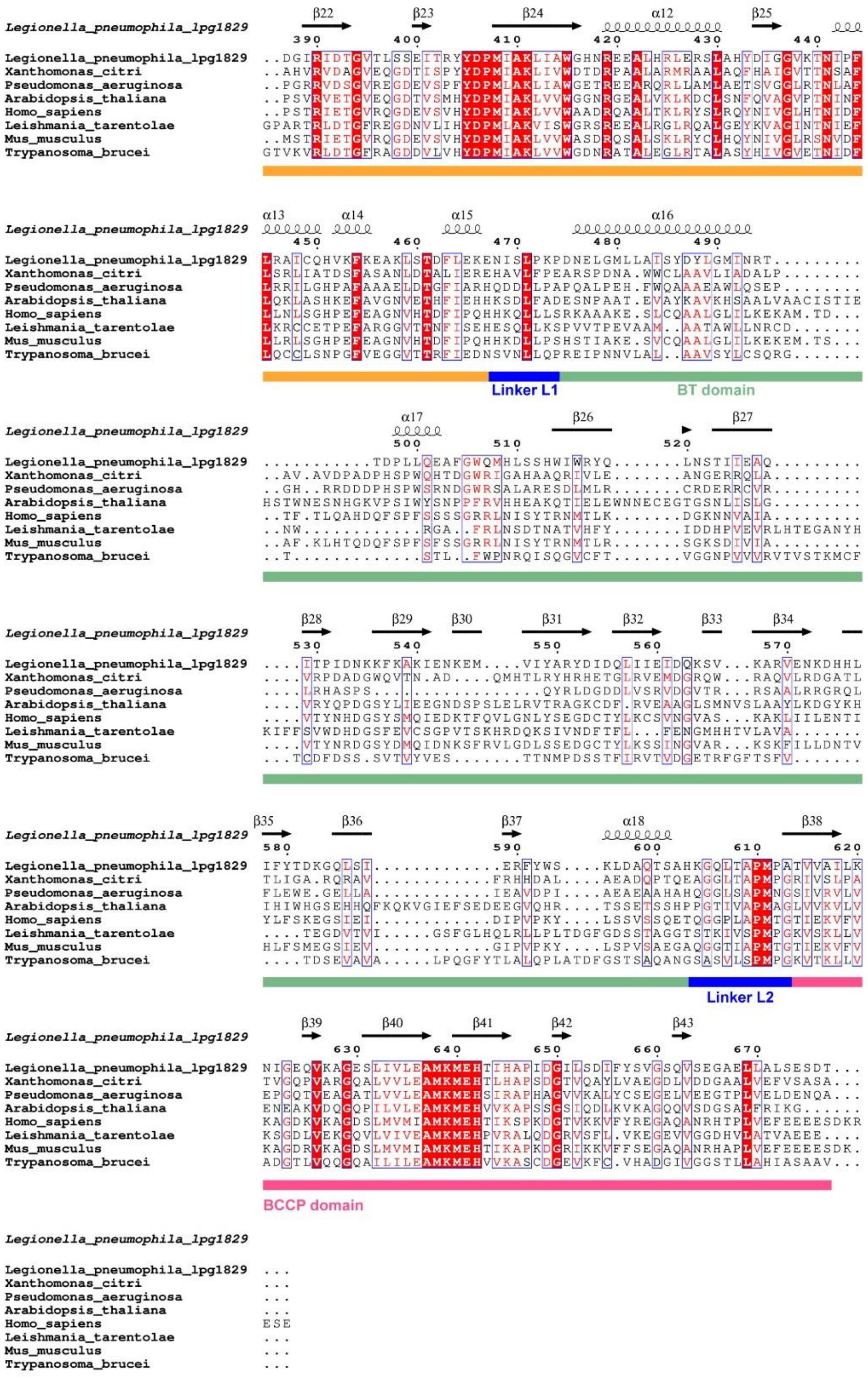

#### Fig. S5 Sequence conservation of the LpMCC α subunit

Multiple sequence alignment of the α subunit from LpMCC and representative bacterial and eukaryotic homologs. Conserved residues are highlighted in red. Secondary structure elements, derived from the LpMCC structure, are shown above the alignment, along with domain boundaries.

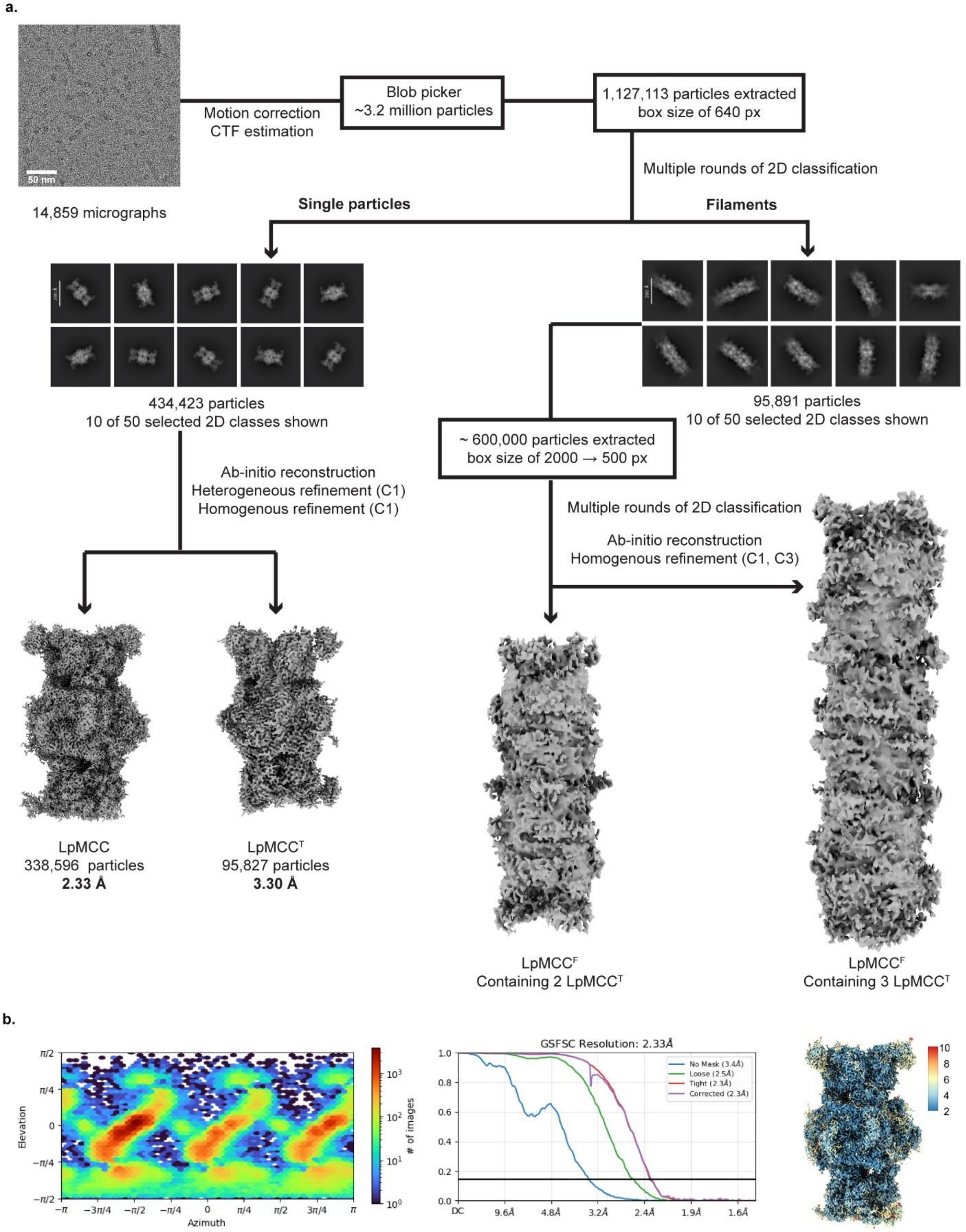
a.
Motion correction
CTF estimation
14,859 micrographs
Blob picker
~3.2 million particles
1,127,113 particles extracted
box size of 640 px
Multiple rounds of 2D classification
Single particles
Filaments
434,423 particles
10 of 50 selected 2D classes shown
95,891 particles
10 of 50 selected 2D classes shown
Ab-initio reconstruction
Heterogeneous refinement (C1)
Homogenous refinement (C1)
~ 600,000 particles extracted
box size of 2000 → 500 px
Multiple rounds of 2D classification
Ab-initio reconstruction
Homogenous refinement (C1, C3)
LpMCC
338,596 particles
2.33 Å
LpMCC^T
95,827 particles
3.30 Å
LpMCC^F
Containing 2 LpMCC^T
LpMCC^F
Containing 3 LpMCC^T
b.
Elevation
Azimuth
### of images
GSFSC Resolution: 2.33Å
No Mask (3.4Å)
Loose (2.5Å)
Tight (2.3Å)
Corrected (2.3Å)
10
8
6
4
2

**Fig. S6 Cryo-EM analysis of substrate-bound LpMCC**

**(a)** Cryo-EM reconstructions of LpMCC in the presence of MCoA, including single-particle states (LpMCC and $LpMCC^{T}$) and filamentous assemblies ($LpMCC^{F}$) containing two or three $LpMCC^{T}$ units. **(b)** Euler angle distribution and gold-standard Fourier shell correlation (GSFSC) curve for substrate-bound LpMCC, with resolution reported at the FSC = 0.143 criterion, and local resolution map of the refined density, colored according to the resolution scale (Å).

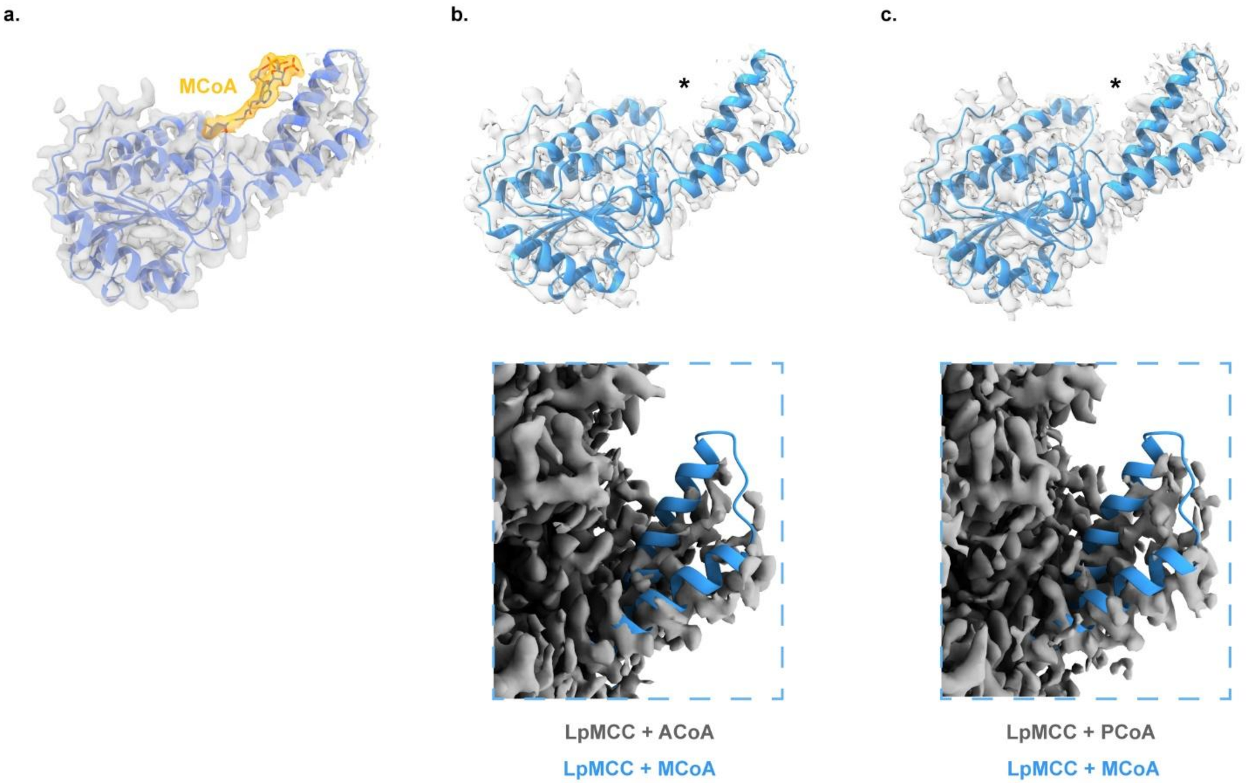


**Fig. S7 CT-C domain density in substrate-bound LpMCC**

**(a)** Model of the CT-C domain from the MCoA–bound LpMCC structure (PDB 9TQ5), with the MCoA substrate shown in orange **(b,c)** Structural superposition of the CT-C domain from the MCoA-bound state (blue) with the ACoA–bound **(b)** and PCoA–bound **(c)** states. The asterisk (*) marks the region where substrate-associated density is observed in the MCoA-bound map but is absent in the ACoA- and PCoA-bound reconstructions. Bottom panels show corresponding cryo-EM density maps, highlighting conformational changes at the helix-turn-helix (HtH) in the presence of ligand density in the MCoA-bound state and its absence in the ACoA- and PCoA-bound states.

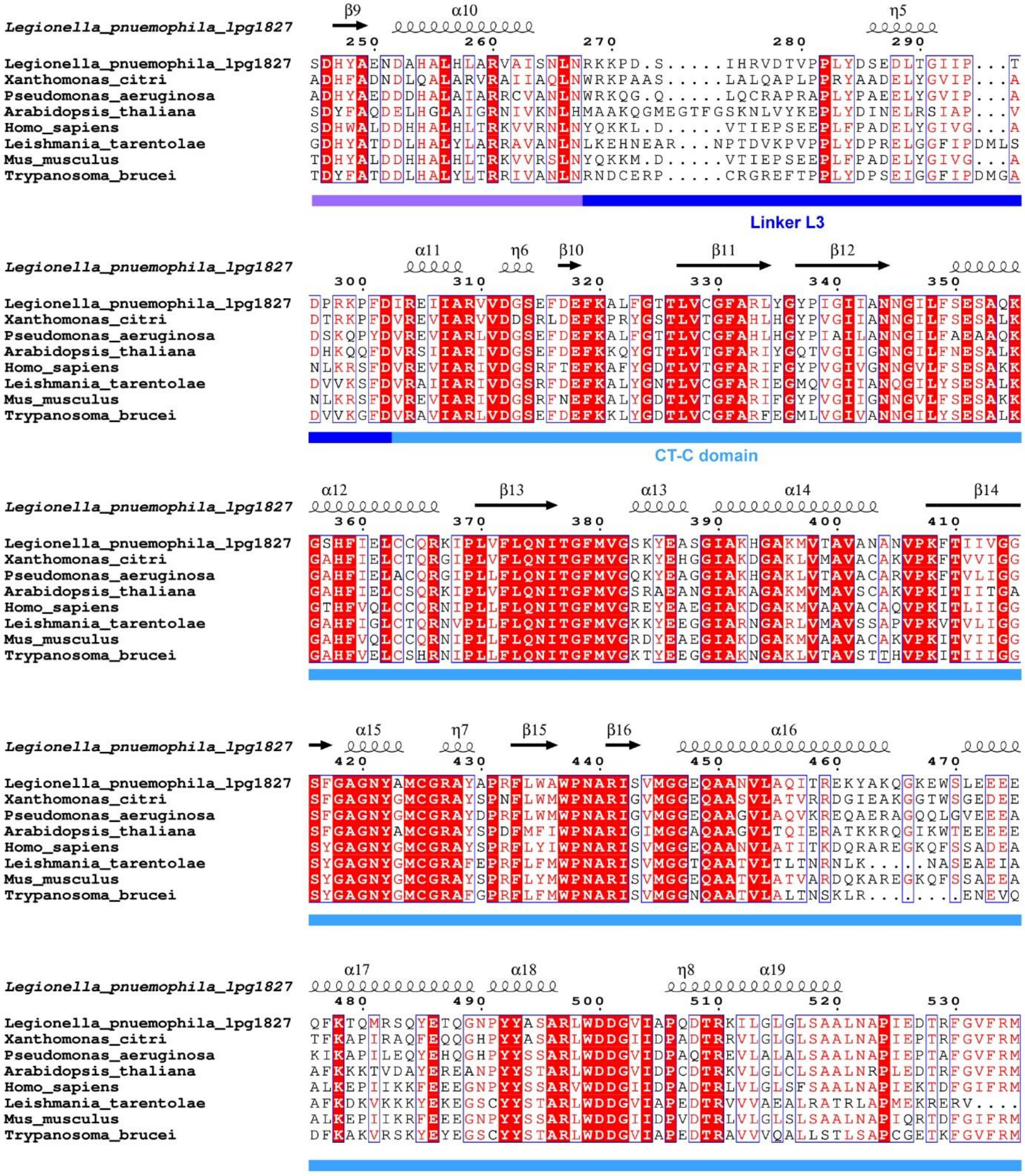


**Fig. S8 Sequence conservation of the LpMCC β subunit.**

Multiple sequence alignment of the β subunit from LpMCC and representative homologs. Conserved residues are highlighted in red. Secondary structure elements, derived from the LpMCC structure, are shown above the alignment, and the CT domain boundaries are indicated.

**Table S1 Cryo-EM data collection, refinement, and validation statistics for LpMCC.**

| | LpMCC | | | With substrate | | |
|---|---|---|---|---|---|---|
| | $\alpha_6\beta_6$ (PDB 9TPS) (EMD-56113) | $\alpha_1\beta_6$ (PDB 9TQG) (EMD-56149) | $\alpha_3\beta_6$ (PDB 9TQC) (EMD-56131) | MCoA (PDB 9TQ5) (EMD-56125) | ACoA | PCoA |
| ***Data collection and processing*** | | | | | | |
| Magnification | 165,000 | 165,000 | 165,000 | 165,000 | 165,000 | 165,000 |
| Voltage (kV) | 300 | 300 | 300 | 300 | 300 | 300 |
| Electron exposure | 40 | 40 | 40 | 40 | 40 | 40 |
| Defocus range (µm) | −0.8 to −1.6 | −0.8 to −1.6 | −0.8 to −1.6 | −0.8 to −1.6 | −0.8 to −1.6 | −0.8 to −1.6 |
| Pixel size (Å) | 0.75 | 0.75 | 0.75 | 0.75 | 0.75 | 0.75 |
| Micrographs collected | 31,641 | 31,641 | 31,641 | 14,859 | 9,011 | 11,825 |
| Resolution (Å) | 2.13 | 2.68 | 2.48 | 2.33 | 2.57 | 2.39 |
| ***Refinement*** | | | | | | |
| Model source | ModelAngelo | PDB 9TPS | PDB 9TPS | PDB 9TPS | | |
| B factor | -41.1 | -41.7 | -38.8 | -30.0 | -46.7 | -48.1 |
| Model composition (Non-H atoms) | 52924 | 25678 | 40132 | 51667 | | |
| Model composition (Protein residues) | 6774 | 3321 | 5148 | 6547 | | |
| Model composition (Ligands) | BTI:6 | BTI:3 | BTI:6 | BTI: 6<br>TW3: 6 | | |
| ***Validation*** | | | | | | |
| MolProbity score | 1.14 | 1.36 | 1.00 | 1.12 | | |
| Clashscore | 2.15 | 2.87 | 1.63 | 1.55 | | |
| Rotamer outliers (%) | 0.84 | 1.42 | 0.36 | 0.8 | | |
| Ramachandran plot (favoured) | 97.13 | 97.0 | 97.58 | 96.6 | | |

| | | | | |
|---|---|---|---|---|
| Ramachandran plot (allowed) | 2.75 | 2.97 | 2.36 | 3.34 |
| Ramachandran plot (Disallowed) | 0.12 | 0.03 | 0.06 | 0.05 |

**Table S2 Protein identification by mass spectrometry (provided as a separate MS Excel file)**
